# Swim exercise in *C. elegans* extends neuromuscular and intestinal healthspan, enhances learning ability, and protects against neurodegeneration

**DOI:** 10.1101/633776

**Authors:** Ricardo Laranjeiro, Girish Harinath, Jennifer E. Hewitt, Jessica H. Hartman, Mary Anne Royal, Joel N. Meyer, Siva A. Vanapalli, Monica Driscoll

## Abstract

Exercise can protect against cardiovascular disease, neurodegenerative disease, diabetes, cancer, and age-associated declines in muscle, immune, and cognitive function. In fact, regular physical exercise is the most powerful intervention known to enhance robustness of health and aging. Still, the molecular and cellular mechanisms that mediate system-wide exercise benefits remain poorly understood, especially as applies to “off target” tissues that do not participate directly in training activity. Elaborating molecular mechanisms of whole-animal exercise benefits is therefore of considerable importance to human health. The development of exercise protocols for short-lived genetic models holds great potential for deciphering fundamental mechanisms of exercise trans-tissue signaling during the entire aging process. Here, we report on the optimization of a long-term swim exercise protocol for *C. elegans* and we demonstrate its benefits to diverse aging tissues, even if exercise occurs only during a restricted phase during early adulthood. We found that multiple daily swim sessions are essential for exercise adaptation in *C. elegans*, leading to body wall muscle improvements in structural gene expression, locomotory performance, and mitochondrial morphology. Swim exercise training enhances whole-animal health parameters such as mitochondrial respiration and mid-life survival and increases the functional healthspan of pharynx and intestine. Importantly, we show that swim exercise also enhances nervous system health: exercise increases learning ability of adult animals and protects against neurodegeneration in *C. elegans* models of tauopathy, Alzheimer’s disease, and Huntington’s disease. An important point is that swim training only during *C. elegans* early adulthood induces long-lasting systemic benefits that in several cases are still detectable well into mid-life. Overall, our data reveal the broad impact of swim exercise in promoting extended healthspan of multiple *C. elegans* tissues, underscore the potency of early exercise experience to influence long-term health (even after cessation of exercise), and establish the foundation for exploiting the powerful advantages of this genetic model to dissect the exercise-dependent molecular circuitry that confers long-lasting system-wide health benefits to aging or diseased adults.

## Introduction

Physical inactivity is a worldwide public health problem. Globally, more than 5 million premature deaths each year (approximately 9%) may be attributable to insufficient physical activity [1]. Conversely, it is striking that exercise provides widespread health benefits, including protection against cardiovascular disease, diabetes, cancer, neurodegenerative disease, and against age-associated declines in muscle, immune, and cognitive function [2, 3]. Given the overwhelming projected social and economic costs of caring for the growing inactive and elderly populations, defining how activity and exercise physiology promote healthier aging is of critical importance. Unfortunately, however, the molecular and cellular mechanisms that mediate systemic exercise benefits remain poorly understood. This gap in knowledge is partially due to the lack of established short-lived genetic models in which fundamental questions on trans-tissue exercise signaling can be evaluated over the entire aging process. Importantly, exercise training in *Drosophila* improves motility, cardiac performance, and mitochondrial health [4–6], supporting that exercise adaptation is conserved from invertebrates to humans, and opening the door to the exploitation of powerful invertebrate model systems for mechanistic dissection of health benefit circuits in exercise biology.

The nematode *Caenorhabditis elegans* is a genetic model widely used in aging and stress research that holds great potential to unlock the molecular circuitry involved in exercise-dependent systemic health benefits. The short lifespan (about 3 weeks) makes *C. elegans* a desirable model to assess exercise effects over the entire adult lifetime. *C. elegans* reach adulthood in ~2.5 days and self-reproduce for approximately the first 6 days of adult life [7]. The reduction in reproductive capacity is followed by age-related decline in multiple tissues and functions over the next several days, including locomotion, muscle structure, pharyngeal pumping, intestinal integrity, and neuronally-controlled actions [7–13]. The aging *C. elegans* body wall muscle exhibits nuclear fragmentation, mitochondrial network and sarcomere disorganization, lipid accumulation, and loss of muscle mass [8, 14], resembling the fundamental characteristics of human sarcopenia (the pervasive age-associated loss of muscle strength and muscle mass that contributes to human frailty).

We have shown previously that swimming in a liquid environment is more energetically demanding for *C. elegans* than the crawling motion on agar plates used in standard lab growth protocols [15]. Moreover, we showed that a single 90 min *C. elegans* swim session induces key features of mammalian exercise, namely locomotory fatigue, muscle mitochondrial oxidation, a transcriptional oxidative stress response, and changes in carbohydrate and fat metabolism [15]. Hartman et al. [16] found that a 6-day swim exercise regimen in *C. elegans* improves body wall muscle mitochondrial morphology, changes mitochondrial respiration parameters, and protects against acute exposure to the mito-toxicants rotenone and arsenic. Other exercise paradigms in *C. elegans* have been described [17–20], with a focus on body wall muscle- and mitochondrial-related phenotypes. However, exercise-dependent effects on a wider range of *C. elegans* tissues, i.e. “off target” benefits of exercise to non-muscle tissue, remain mostly undescribed.

Here, we report on the optimization of a long-term swim exercise protocol for *C. elegans* and establish that multiple daily swim sessions are essential for exercise adaptation. Body wall muscle directly benefits from swim exercise training and features muscle structural gene expression increase, improved locomotory performance, and enhanced maintenance of the muscle mitochondrial network. Whole-animal parameters such as mitochondrial respiration and mid-life survival are also improved in exercised *C. elegans* as compared to non-exercised controls. Importantly, we show that swim exercise improves pharyngeal, intestinal, and neuronal functions. Furthermore, we find that neurodegenerative pathological conditions in *C. elegans* models of tauopathy, Alzheimer’s disease (AD), and Huntington’s disease (HD) are ameliorated by swim exercise training. The systemic exercise effects we describe reveal a broad impact of swim training on *C. elegans* physiology and establish the critical groundwork for future dissection of the molecular circuitry responsible for exercise-dependent trans-tissue health benefits.

## Results

### Multiple daily swim sessions are essential for *C. elegans* exercise adaptation

Although a single 90 min swim session in *C. elegans* induces key features of mammalian exercise [15], most exercise-dependent health benefits in humans arise from long-term training regimens that lead to exercise adaptation over time [21]; thus, we sought to develop and validate a long-term *C. elegans* exercise protocol. We planned to determine an efficacious long-term exercise regimen in *C. elegans* by characterizing the impact of different numbers of 90 min swim sessions in M9 buffer administered over the first four days of adult life (Fig. 1a). The first days of adult life correspond to the peak locomotory performance phase and constitute a timeframe over which *C. elegans* are able to complete 90 min swims without any significant inactive periods [9, 10, 22]. We tested long-term exercise regimens with swim frequencies of: one session per day (1+1+1+1); two sessions per day (2+2+2+2); three sessions per day on the first two days of adulthood plus two sessions per day on the third and fourth adult days (3+3+2+2); three sessions per day (3+3+3+3); and four sessions per day (4+4+4+4). The exercise adaptation readouts we quantitated included both a molecular assay (gene expression analysis by quantitative PCR (qPCR)) and a functional assay (locomotion analysis by recording movement and calculating crawling maximum velocity). Importantly, given that our interest is in defining regimens that lead to medium-to long-term physiological changes (i.e., exercise adaptation and system-wide health benefits), we assayed molecular and functional readouts one to several days post-exercise, rather than immediately after the last swim session.

**Figure 1.**
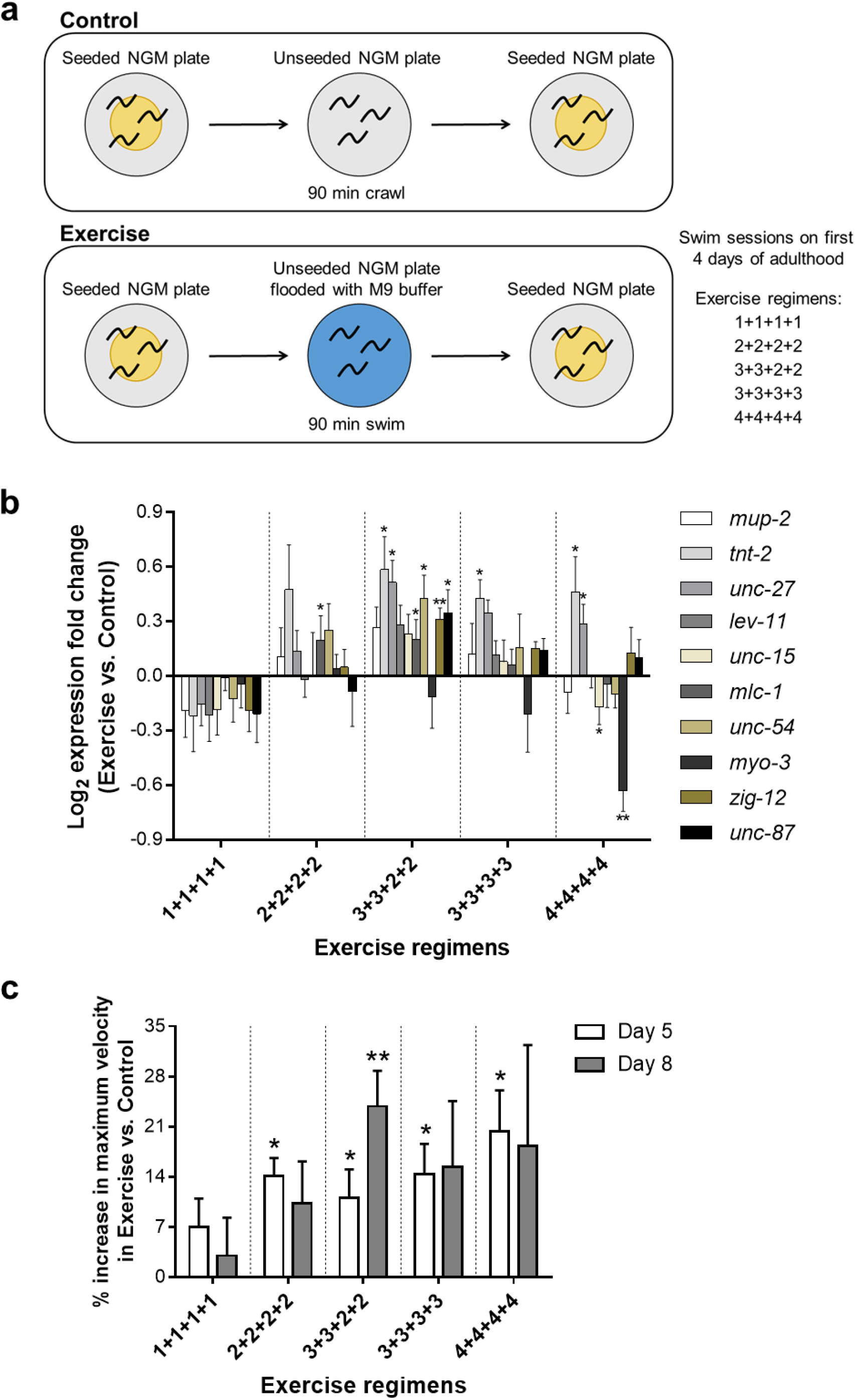
Multiple daily swim sessions are essential for *C. elegans* exercise adaptation. **a**, Diagram of the long-term swim exercise regimens tested. We transferred control animals to an unseeded NGM plate for 90 min, whereas exercise animals were transferred to an unseeded NGM plate flooded with M9 buffer for the same 90 min to swim. This experimental design controls for any differences between control and exercise nematodes that might be attributed to differences in food availability or animal handling. Swim frequency on successive days is indicated. For example, 3+3+2+2 indicates 90 min swims 3X per day on Ad1, Ad2 and 2X per day on Ad3, Ad4. **b**, qPCR results for 10 genes encoding proteins important for muscle contraction in Ad5 WT animals exposed to the five long-term exercise regimens indicated. Expression data are presented as the log_2_ fold change of exercise samples relative to control samples. Number of independent trials performed: *n*_1+1+1+1_ = 4, *n*_2+2+2+2_ = 4, *n*_3+3+2+2_ = 5, *n*_3+3+3+3_ = 4, *n*_4+4+4+4_ = 4. We used 30-40 animals per sample in each trial. Statistical significance determined by paired two-tailed Student’s *t* test. **c**, Percentage increase in crawling maximum velocity in Ad5 and Ad8 WT exercised animals, exposed to the five tested long-term exercise regimens, relative to non-exercised control counterparts. Number of independent trials performed: *n*_Ad5 1+1+1+1_ = 4, *n*_Ad8 1+1+1+1_ = 4, *n*_Ad5 2+2+2+2_ = 3, *n*_Ad8 2+2+2+2_ = 3, *n*_Ad5 3+3+2+2_ = 5, *n*_Ad8 3+3+2+2_ = 6, *n*_Ad5 3+3+3+3_ = 4, *n*_Ad8 3+3+3+3_ = 4, *n*_Ad5 4+4+4+4_ = 3, *n*_Ad8 4+4+4+4_ = 3. We used ~20 animals per sample in each trial. Statistical significance determined by paired two-tailed Student’s *t* test. Data are mean ± s.e.m. \**P*<0.05, \*\**P*<0.01.

Our previous analysis of single swim sessions supported that swimming is an endurance-like exercise for *C. elegans* [15]. In mammals, long-term endurance exercise leads to profound transcriptional changes at the muscle level, with upregulation of muscle structural genes as one of the reproducible changes [23–25]. Therefore, we first wanted to determine if long-term swim exercise regimens could also promote the upregulation of muscle structural genes in *C. elegans*. We selected 10 genes encoding proteins important for muscle contraction, including troponins, tropomyosin, myosins, and titin [26] (Table S1). Importantly, all the genes we included in our analysis have been documented to be massively downregulated with age in *C. elegans* [26], which both underscores their potential importance in age-associated sarcopenia and locomotory decline and increases the potential to detect differential transcriptional changes after long-term exercise regimens. We performed qPCR for the test swim regimens one day after the training was completed, which corresponds to Adult day 5 (Ad5) (Fig. 1b). We find that the 1+1+1+1 regimen led to an overall tendency of downregulation across the muscle structural gene test set, supporting that single daily swim sessions are insufficient to induce exercise adaptations (at least at the molecular level). By contrast, 2+2+2+2, 3+3+2+2, and 3+3+3+3 regimens led to an overall upregulation of muscle structural gene expression. The 3+3+2+2 regimen induced the most consistent transcriptional upregulation profile with 6 out of the 10 test genes (*tnt-2*, *unc-27*, *mlc-1*, *unc-54*, *zig-12*, and *unc-87*) exhibiting a statistically significant difference when compared to the control animals. The 4+4+4+4 regimen induced a wider range of transcriptional outcomes (two genes showing a significant upregulation (*tnt-2* and *unc-27*) and two genes showing a significant downregulation (*unc-15* and *myo-3*)) and reflected a trend toward downregulation that suggested excessive exercise can be deleterious to maintained muscle expression. Overall, our data establish that training plans of 2+2+2+2, 3+3+2+2, and 3+3+3+3 can induce increased transcription of a battery of *C. elegans* muscle structural genes, paralleling a feature of mammalian exercise adaptation.

To assess training impact on muscle performance, we also measured a functional readout of our long-term *C. elegans* exercise regimens. We focused on the *C. elegans* maximum velocity measure as a powerful functional readout because maximum velocity has been documented to accurately report movement ability, correlates well with healthspan, and at certain ages can even be a longevity predictor [10]. We measured maximum velocity by placing Ad5 and Ad8 single animals on agar plates and determining the maximum speed at which they move over a 30-second time interval. Note that our swim-exercised animals have not been specifically trained for enhancement of crawling maximum velocity, so maximum velocity serves as an indicator of overall vigor that results from swim training. We find that the 1+1+1+1 regimen failed to significantly increase maximum velocity of exercised animals compared to controls on both days (Fig. 1c), independently supporting our conclusion from transcriptional studies that single daily swim sessions are not sufficient to induce long-term exercise adaptations. In contrast, we found that all the multiple daily swim regimens promoted a significant increase in maximum velocity, on the order of a 10-20% increase over non-exercised animals assayed at Ad5, with the 4+4+4+4 regimen showing the best improvement (Fig. 1c). We found a trend toward increased maximum velocity on Ad8 (4 days after cessation of training) for all four multiple daily swim regimens, but only the 3+3+2+2 regimen led to a statistically significant 24% increase (Fig. 1c). Our findings reveal a remarkable long-term benefit of sustained early adult life exercise training.

Overall, the molecular and functional readouts we tested establish that a single 90 min swim session per day does not suffice to promote exercise adaptation in *C. elegans*. Rather, multiple daily swim sessions are required in wild type (WT) *C. elegans* for robust exercise adaptation at molecular and functional levels, with the 3+3+2+2 regimen inducing improvements in multiple assays and with long-term functional maintenance.

### Molecular and functional changes are dependent on increased locomotor activity rather than exposure to a liquid environment

We showed previously that a single swim session in M9 buffer induces specific exercise-dependent acute responses and does not activate a generalized stress response due to liquid exposure [15]. Nevertheless, it remained possible that continuous re-exposure of *C. elegans* to the M9 buffer liquid environment during the long-term exercise regimens might induce some physiological changes. To definitively rule out simple liquid exposure as an important factor for the molecular and functional changes we documented above, we designed microfluidic devices in which *C. elegans* can be exposed to M9 buffer but in which we can manipulate the locomotion type (crawling vs. swimming) by modulating the chamber height. We based the design of our microfluidic test devices on the findings by Lebois et al. that *C. elegans* locomotion can be modulated through confinement [27]. When the microfluidic chamber height is larger than the worm diameter, animals are able to swim (Video S1a), whereas when the microfluidic chamber height is smaller than the worm diameter, animals are physically confined and adopt a slow crawling motion (Video S1b). To test whether exposure to the M9 buffer swim media might suffice to induce exercise adaptations, we performed the 3+3+2+2 regimen by transferring *C. elegans* from seeded nematode growth medium (NGM) plates to M9 buffer-filled microfluidic devices that had a large chamber height (swim exercise possible) or a small chamber height (control, animals are sedate and crawl rather than swim) for 90 min. After each 90 min session, we recovered animals from the microfluidic chambers and returned them to seeded NGM plates. We conducted qPCR analysis at Ad5 to find that microfluidics-exercised *C. elegans* upregulate muscle structural genes (statistically significant for *tnt-2*, *mlc-1*, *unc-54*, and *zig-12*) as compared to microfluidics-control animals (Fig. 2a), similar to the outcomes of the 3+3+2+2 regimen performed on NGM plates (Fig. 1b). Additionally, we find that the microfluidics-exercised animals exhibited a significant increase in maximum velocity on Ad8 relative to microfluidics non-exercised animals, similar to the 3+3+2+2 regimen implemented by flooding plates with M9 buffer (Fig. 2b). Our data establish that transcriptional and functional changes induced in M9 buffer require swim activity and are not the consequence of liquid exposure. Data also independently confirm that long-term swim regimens induce exercise adaptations.

**Figure 2.**
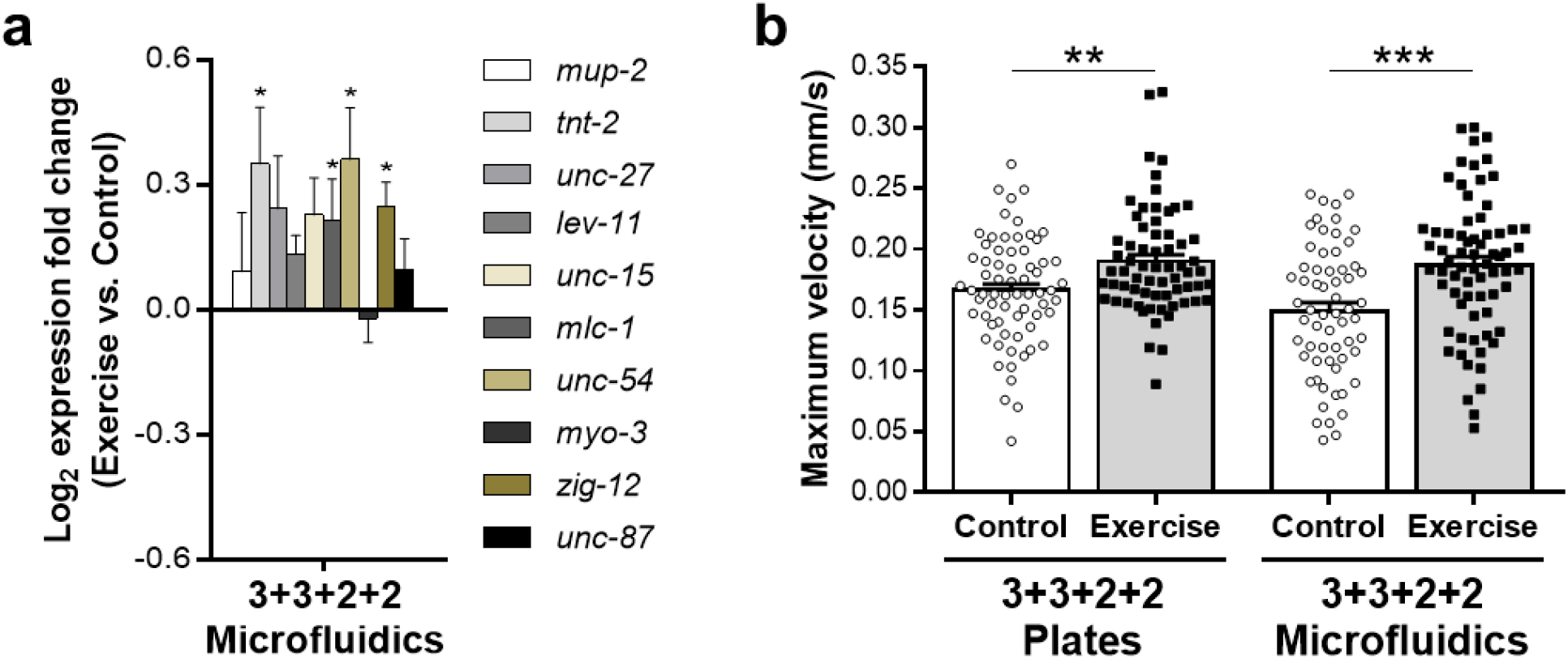
Exercise adaptations depend on increased locomotor activity rather than exposure to a liquid environment. **a**, qPCR results for 10 genes encoding proteins important for muscle contraction in Ad5 WT animals that executed the 3+3+2+2 regimen in microfluidic devices. Expression data are presented as the log_2_ fold change of exercise samples (microfluidic device permissive for swimming) relative to control samples (microfluidic device that restricts locomotion to prevent swimming in the liquid environment of the chamber). *n* = 4 independent trials. We used ~20 animals per sample in each trial. Statistical significance determined by paired two-tailed Student’s *t* test. **b**, Maximum velocity of Ad8 WT animals exposed to the 3+3+2+2 regimen on NGM plates (left) and in microfluidic devices (right). Each point represents a single animal. Number of animals used for analysis: *n*_Control Plates_ = 67, *n*_Exercise Plates_ = 66, *n*_Control Microfluidics_ = 62, *n*_Exercise Microfluidics_ = 70. Data compiled from 4 independent trials. Statistical significance determined by unpaired two-tailed Student’s *t* test. Data are mean ± s.e.m. \**P*<0.05, \*\**P*<0.01, \*\*\**P*<0.001.

### Long-term swim exercise improves performance and healthspan of *C. elegans* body wall muscle

The *C. elegans* body wall muscle is the tissue most directly involved in long-term exercise training and thus we sought to characterize additional aspects of body wall muscle performance and health consequent to exercise. We recently developed a novel burrowing assay, in which nematodes are stimulated to move through a Pluronic F-127 gel toward a chemoattractant [28]. Pluronic F-127 is a hydrogel that transitions from liquid to gel at safe temperatures for *C. elegans*. We loaded liquid-phase Pluronic F-127 on top of *C. elegans* in 12-well plates, and allowed the Pluronic F-127 to quickly gel at room temperature, trapping animals under the gel. We then added *E. coli* OP50-1 on top of the gel (Fig. 3a). In this 3D gel environment, animals with a better neuromuscular performance are able to reach the surface faster and in higher numbers [28]. After a 3+3+2+2 training regimen, Ad5 exercised nematodes reached the gel surface at a significantly higher proportion during the 3-hour burrowing assay window as compared to their non-exercised control counterparts (log-rank test comparing control vs. exercise curves, *P* = 0.013) (Fig. 3b), an additional line of evidence that long-term swim exercise improves the functional output of *C. elegans* body wall muscle.

**Figure 3.**
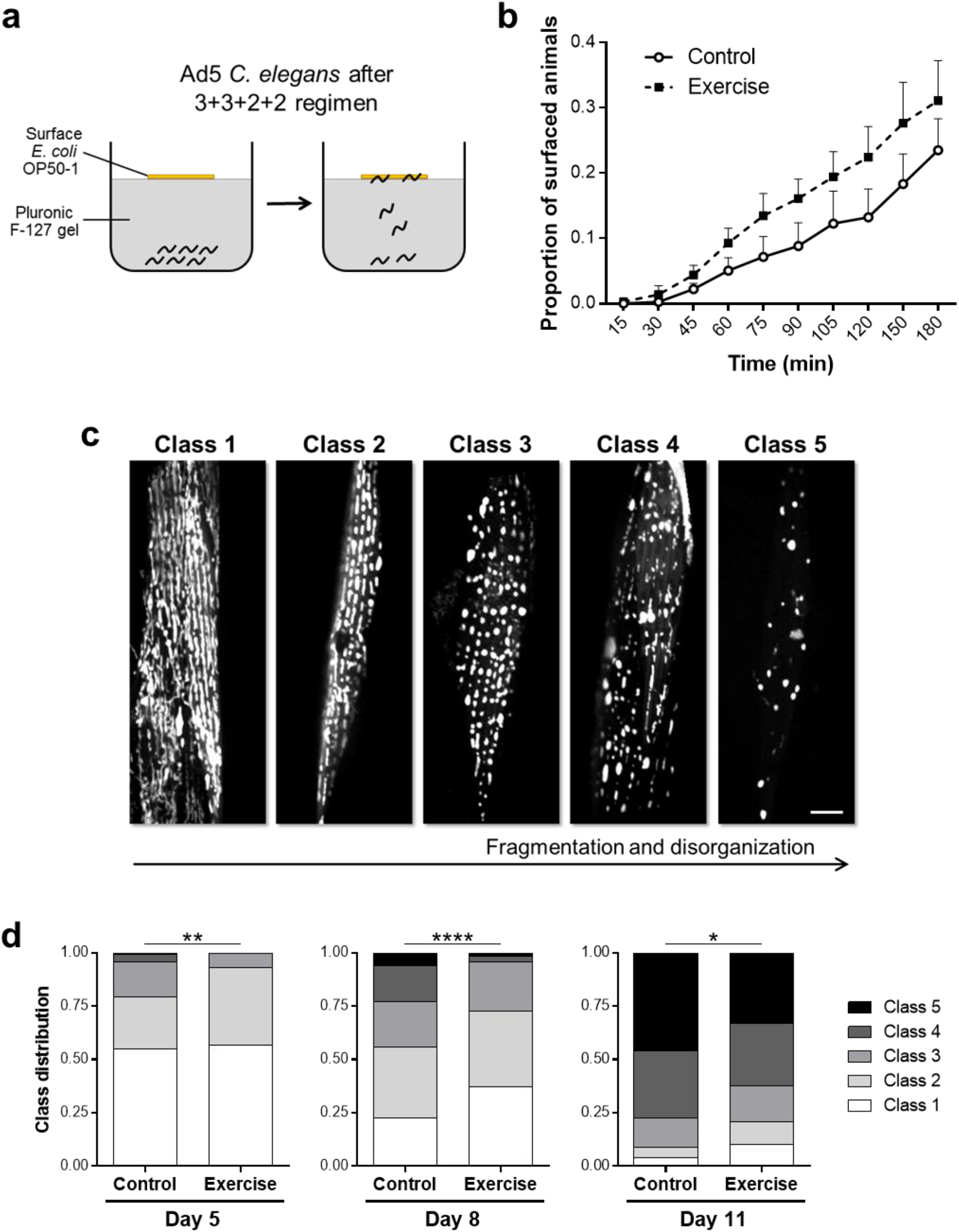
Long-term swim exercise improves burrowing performance and mitochondrial profiles of body wall muscle. **a**, Diagram of burrowing assay. After the 3+3+2+2 regimen, we trapped Ad5 animals under a Pluronic F-127 gel and added attractant food *E. coli* OP50-1 to the center of the gel surface. Animals attracted by the food burrow to the surface of the gel at different rates. **b**, Proportion of Ad5 WT animals exposed to the 3+3+2+2 regimen that reach the gel surface during the 3-hour burrowing assay. Number of animals used for analysis: *n*_Control_ = 392, *n*_Exercise_ = 370. Data compiled from 3 independent trials. Statistical significance determined by log-rank test comparing control vs. exercise curves (*P* = 0.013). **c**, Representative confocal images of the five classes of body wall muscle mitochondrial network organization in *P_myo-3_ mitoGFP* animals. Fragmentation and disorganization of muscle mitochondria increases progressively from Class 1 to Class 5. Scale bar: 10 µm. **d**, Distribution of mitochondrial classes in body wall muscle of Ad5, Ad8, and Ad11 *P_myo-3_mitoGFP* animals exposed to the 3+3+2+2 regimen. We scored the confocal muscle images blind to exercise condition. Number of body wall muscle images used for analysis: *n*_Ad5 Control_ = 170, *n*_Ad5 Exercise_ = 160, *n*_Ad8 Control_ = 160, *n*_Ad8 Exercise_ = 162, *n*_Ad11 Control_ = 160, *n*_Ad11 Exercise_ = 160. Data compiled from 4 independent trials. Statistical significance determined by chi-square test. Data are mean ± s.e.m. \**P*<0.05, \*\**P*<0.01, \*\*\*\**P*<0.0001.

One of the hallmarks of endurance exercise in mammals is muscle mitochondrial adaptation, which can be reflected in morphological and functional mitochondrial improvements [29]. We therefore explored the consequences of long-term swim exercise on the body wall muscle mitochondrial network by using a *C. elegans* strain in which mitochondria are labeled with GFP specifically in the body wall muscle (*P_myo-3_mitoGFP*). *C. elegans*, just like mammals, exhibit a well-described age-related decline in muscle mitochondrial morphology, characterized by increased disorganization and fragmentation of the mitochondrial network [14, 26, 30–32]. We performed confocal imaging of *P_myo-3_mitoGFP* animals at different life stages to establish our own classification system for muscle mitochondrial network health that reflects a progressive increase in fragmentation and disorganization from Class 1 (complete mitochondrial coverage of body wall muscle cells and tubular mitochondrial morphology) to Class 5 (greatly reduced number of mitochondria and round morphology) (Fig. 3c). After the 3+3+2+2 exercise regimen, we took confocal images of the anterior body wall muscle of *P_myo-3_mitoGFP* animals on Ad5, Ad8, and Ad11, and we scored the images blind to exercise condition using the freely-available software Blinder [33]. We find that exercised animals exhibited a significant improvement in muscle mitochondrial network organization at all three time points tested (Fig. 3d), even though Ad11 corresponds to a full 7 days after the last exercise session. Our results show that long-term swim exercise in *C. elegans* promotes long-lasting changes in the body wall muscle mitochondrial populations that are correlated with improvements in muscle performance and healthspan.

### Long-term exercise improves whole-animal mitochondrial respiration parameters

A central question in exercise physiology is the molecular nature of the factors that promote systemic health consequent to exercise. Given that the *C. elegans* model holds potential for genetic dissection of such health signaling circuits, we turned to focus on assessing animal-wide consequences of exercise for aging adults, with the expectation of defining endpoints that could be used for future mechanistic dissection.

In light of the documented benefits of long-term swim exercise to muscle mitochondrial networks, we first sought to determine if mitochondrial function as measured at the whole-animal level was affected by our exercise regimens. To address this question, we analyzed numerous parameters of mitochondrial respiration in whole nematodes by measuring oxygen consumption rates with a Seahorse XFe24 Analyzer. In the Seahorse assays for measuring oxygen consumption rates, basal respiration is first monitored; dicyclohexylcarbodiimide (DCCD) is then added to inhibit ATP synthase, which reveals relative ATP-linked respiration and distinguishes the residual proton leak; carbonyl cyanide-p-trifluoromethoxyphenylhydrazone (FCCP) then uncouples mitochondria, collapsing the proton gradient such that the electron transport chain (ETC) is uninhibited and oxygen consumption by complex IV reaches the maximal rate (the difference between maximal respiration and basal respiration is the spare capacity); addition of sodium azide (a cytochrome c oxidase inhibitor) shuts down all mitochondrial respiration to reveal non-mitochondrial respiration [34] (Fig. 4a).

**Figure 4.**
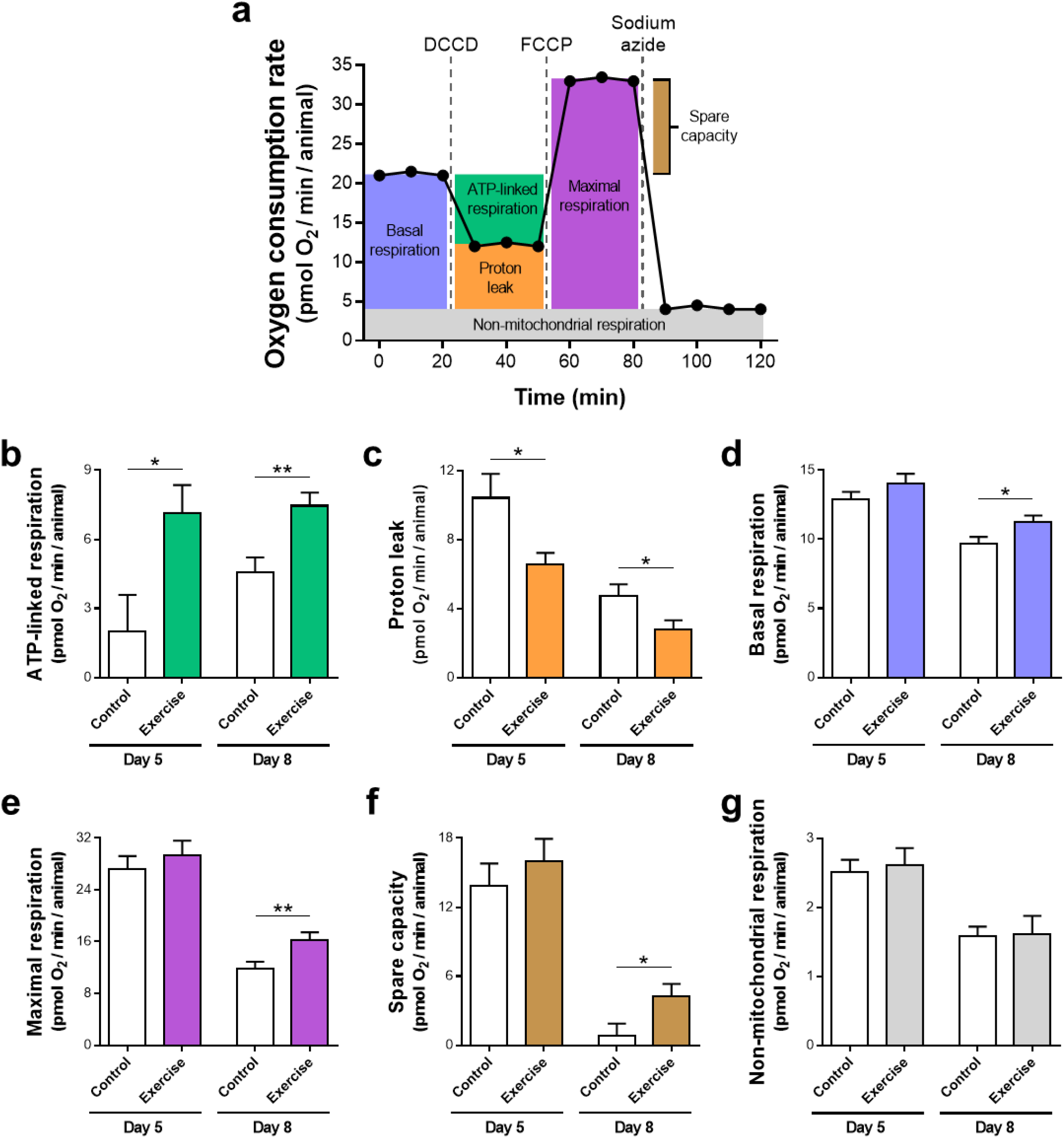
Long-term exercise improves whole-animal mitochondrial respiration parameters. **a**, Diagram of oxygen consumption rates measured in a Seahorse XFe24 Analyzer consequent to addition of mitochondrial inhibitors. In the Seahorse protocol for measuring oxygen consumption rates, basal respiration is first monitored followed by addition of DCCD, FCCP, and sodium azide allowing for the calculation of the six respiration parameters represented by different colors. **b-g**, ATP-linked respiration (**b**), proton leak (**c**), basal respiration (**d**), maximal respiration (**e**), spare capacity (**f**), and non-mitochondrial respiration (**g**) values of Ad5 and Ad8 WT animals exposed to the 4+4+4+4 regimen. Number of Seahorse XF24 Microplate wells used for analysis: *n*_Ad5 Control_ = 18 in **b**, **c**, **e**, **f**, and **g**, *n*_Ad5 Control_ = 36 in **d**, *n*_Ad5 Exercise_ = 22 in **b**, **c**, **e**, **f**, and **g**, *n*_Ad5 Exercise_ = 44 in **d**, *n*_Ad8 Control_ = 19 in **b**, **c**, and **g**, *n*_Ad8 Control_ = 39 in **d**, *n*_Ad8 Control_ = 20 in **e** and **f**, *n*_Ad8 Exercise_= 19 in **b**, **c**, and **g**, *n*_Ad8 Exercise_ = 41 in **d**, *n*_Ad8 Exercise_ = 22 in **e** and **f**. Data compiled from 4 independent trials. We used 20-25 animals per well in each trial. Statistical significance determined by unpaired two-tailed Student’s *t* test. Data are mean ± s.e.m. \**P*<0.05, \*\**P*<0.01.

We swim-exercised *C. elegans* with a 4+4+4+4 regimen (a training regimen we used prior to optimization described above) and assayed oxygen consumption rates the following day (Ad5) and four days after training (Ad8). We find that exercised animals exhibited an increase in ATP-linked respiration (Fig. 4b) concomitant with a decrease in proton leak (Fig. 4c), at both Ad5 and Ad8, as compared to non-exercised control counterparts. Since the proton gradient is used preferentially for ATP production in exercised animals, and in non-exercised control animals a large proportion of the proton gradient is lost due to proton leak, we conclude that mitochondrial respiration is more efficient in exercised vs. control animals. We did not record any differences in other parameters measured between exercise and control animals at Ad5. However, at Ad8, basal respiration, maximal respiration, and spare capacity were all significantly increased in exercised animals as compared to controls (Fig. 4d-f). Higher basal and maximal respiration in the exercised cohort suggest more and/or healthier mitochondria in exercised animals. Importantly, the increase in spare capacity at Ad8 reflects an improved ability of exercised animals to meet energy requirements under potential challenges of high-energy demand situations. We observed no differences for non-mitochondrial respiration values between control and exercise samples (Fig. 4g), suggesting that swim exercise training specifically enhances mitochondrial function rather than affecting overall metabolic reactions.

Taken together, our analyses support that long-term swim exercise in *C. elegans* induces both mitochondrial morphological changes (in muscle) and improvements in whole-animal mitochondrial efficiency and health, with effects that can last for several days after exercise cessation.

### Mid-life survival is increased in swim-exercised *C. elegans*

The improvement in mitochondrial function we documented at the whole-animal level raised the question whether swim exercise is able to improve additional organismal-wide health measures. In humans, exercise can be correlated with longevity [35–38], and thus we compared relative survival of 3+3+2+2 exercised and non-exercised populations to address overall health. Our survival assays revealed that a 3+3+2+2 swim exercise regimen does not change the maximum lifespan of *C. elegans* population (Fig. 5). However, exercised animals showed on average ~15% increase in survival between Ad12 and Ad20 compared to control animals (log-rank test comparing control vs. exercise survival curves, *P* = 0.002) (Fig. 5). This enhancement in mid-life survival provides further evidence that long-term swim exercise in *C. elegans* induces systemic health benefits rather than isolated adaptations at the body wall muscle level.

**Figure 5.**
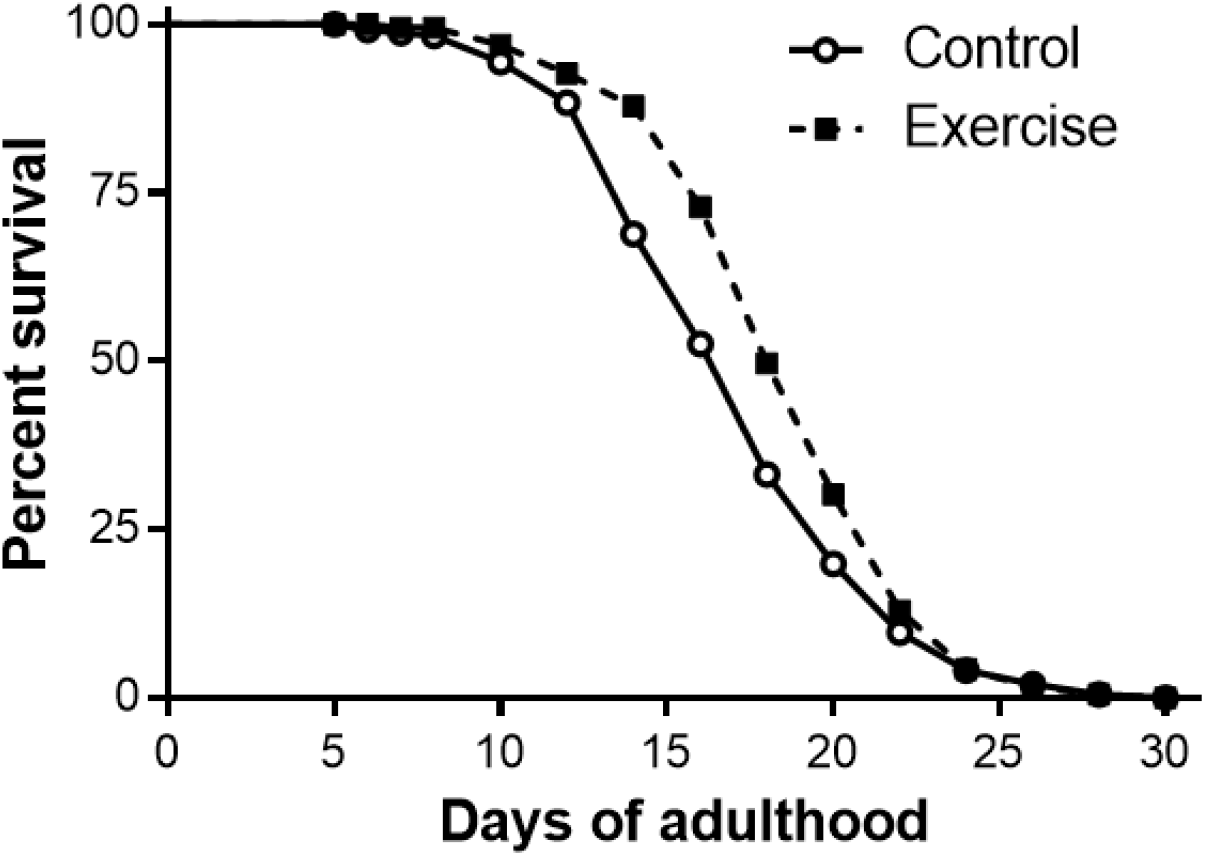
Mid-life survival is increased in swim-exercised *C. elegans*. Survival curve of WT animals exposed to the 3+3+2+2 regimen. Number of animals used for analysis: *n*_Control_ = 245, *n*_Exercise_ = 221. Data compiled from 3 independent trials. Statistical significance determined by log-rank test comparing control vs. exercise survival curves (*P* = 0.002).

### Pharyngeal and intestinal healthspans are extended after long-term swim exercise

Given swim exercise benefits evident at the whole-nematode level, we sought to detect tissue-specific improvements in *C. elegans* organs that are not an activity focus during exercise. The *C. elegans* pharynx is of particular interest as pharyngeal pumping function declines markedly with age [7] and the pharynx shares molecular and functional similarities with the vertebrate heart, including continuous pumping of material along the lumen of a muscle tube over life; electrical circuitry (contractions synchronized by gap junctions that can continue in the absence of neuronal input and are modulated by neurotransmitters); transcription factors involved in biogenesis; and conserved cardiac muscle proteins [39]. Notably, however, *C. elegans* swim exercise does not enhance pumping during swimming [40], so that training benefits are anticipated to be secondary effects of the swim.

We quantified pharyngeal pumping rate of WT *C. elegans* after a 3+3+2+2 exercise regimen. We observed no difference in pumping rate at Ad8 but at Ad11 and Ad15, exercised animals showed a significant increase in pumping rate relative to controls (Fig. 6a). Notably, exercise health benefits to pharyngeal function become evident during the time-window of accentuated age-related decline. Once again, even though animals are trained only on the first four days of adulthood, exercise-dependent effects are long lasting and in this case maintained 11 days after the last exercise bout.

**Figure 6.**
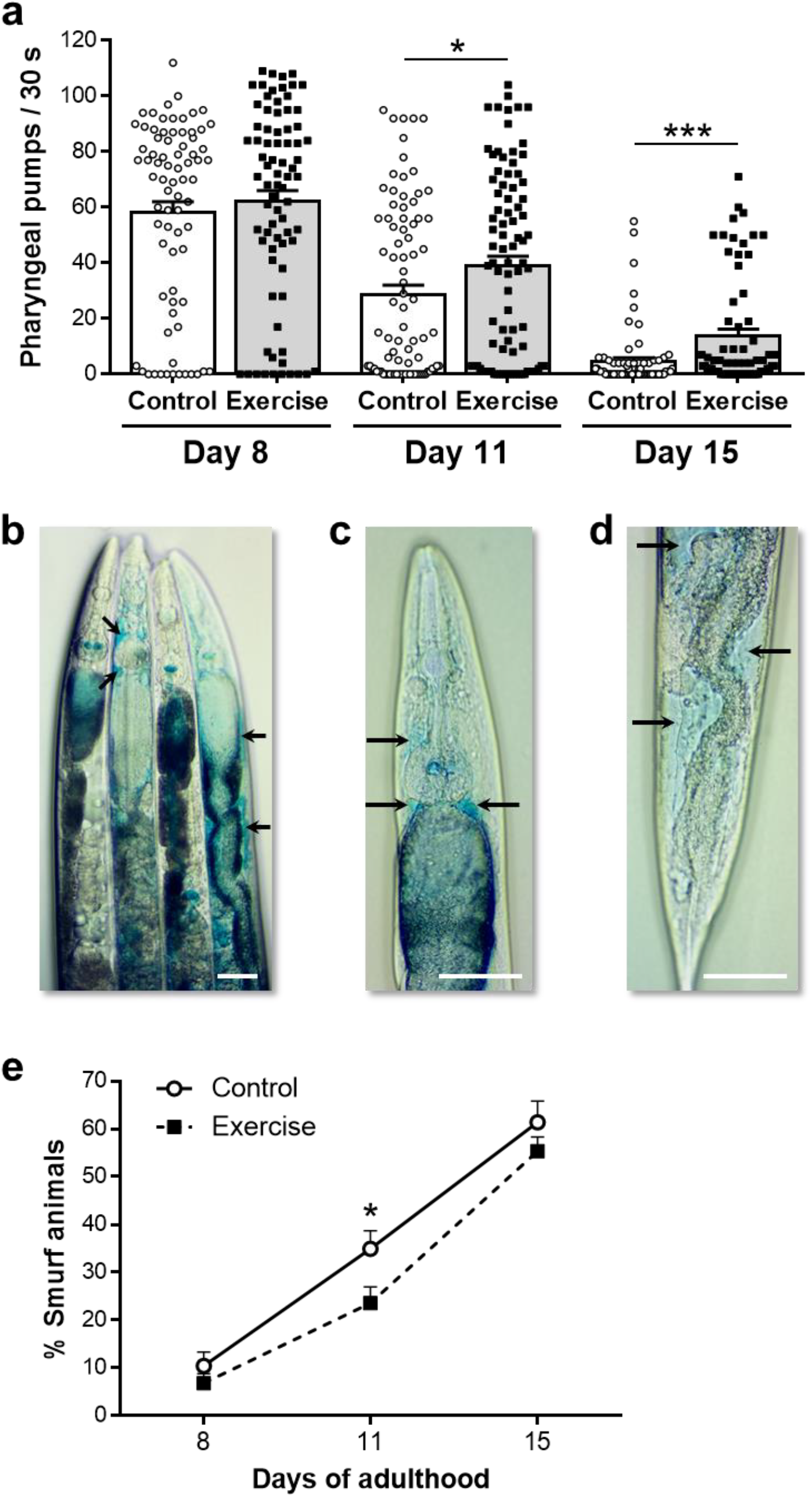
Pharyngeal and intestinal healthspans are extended after long-term swim exercise. **a**, Pharyngeal pumping rate of Ad8, Ad11, and Ad15 WT animals exposed to the 3+3+2+2 regimen. Each point represents a single animal. Number of animals used for analysis: *n*_Ad8 Control_ = 76, *n*_Ad8 Exercise_ = 77, *n*_Ad11 Control_ = 85, *n*_Ad11 Exercise_ = 87, *n*_Ad15 Control_ = 75, *n*_Ad15 Exercise_ = 69. Data compiled from 3 independent trials. Statistical significance determined by unpaired two-tailed Student’s *t* test. **b**, Representative image of non-Smurf (first and third animals from left to right) and Smurf (second and fourth animals from left to right, arrows indicate leaks) animals. **c, d**, Higher magnification images of Smurf animals showing intestinal leakage in the anterior (**c**) and posterior (**d**) regions. Arrows in **b-d** indicate areas of blue dye leakage into the body cavity. Scale bars: 50 µm. **e**, Percentage of Smurf animals at Ad8, Ad11, and Ad15 in WT nematodes exposed to the 3+3+2+2 regimen. Number of independent trials performed: *n*_Ad8_ = 4, *n*_Ad11_ = 8, *n*_Ad15_ = 8. We used 25-40 animals per sample in each trial. Statistical significance determined by paired two-tailed Student’s *t* test. Data are mean ± s.e.m. \**P*<0.05, \*\*\**P*<0.001.

The *C. elegans* intestine is an additional tissue of central health importance, as the intestine plays critical roles in digestion and metabolism and serves as a focal signaling center for stress response and aging regulation [41]. The *C. elegans* intestine constitutes a strong barrier to gut content leak in young animals, but with age, intestinal barrier function breaks down. Breakdown of intestinal barrier function can be measured by feeding a non-absorbable blue dye and checking for dye leakage outside the intestine, an assay referred to as the “Smurf assay” since animals turn blue like cartoon Smurfs [42–44]. We assayed the maintenance of gut integrity in exercised vs. non-exercised controls. We performed the Smurf assay by incubating *C. elegans* in a bacterial liquid culture with blue dye for 3h. Animals with intestinal epithelial integrity retained the blue dye within the intestinal tract, whereas animals with a compromised intestinal barrier leak the blue dye to their body cavity (Smurf animals) (Fig. 6b-d). We find that the percentage of Smurf animals in both control and exercise samples was very low at Ad8 after a 3+3+2+2 regimen (Fig. 6e), consistent with prior studies that revealed the later life onset of gut decline [44]. Later in life (Ad11), we find that the proportion of Smurf animals was one-third lower in exercise versus control (Fig. 6e). Differences disappeared in older animals – at Ad15 more than 50% of both populations exhibited intestinal leakage (Fig. 6e). We conclude that long-term swim exercise improves intestinal integrity and function for a particular time-window later in *C. elegans* life.

### Learning ability is enhanced in exercised *C. elegans*

Benefits to nervous system function, including learning and memory enhancement, are documented outcomes of mammalian exercise [45–48]. We therefore tested whether long-term exercise improves associative neuronal functions in adult *C. elegans*. Positive learning assays in which *C. elegans* associate food with a specific chemical odor (e.g. butanone) have been well described [13, 49]. We tested a paradigm in which animals are starved for 1h followed by food-butanone conditioning in seeded NGM plates with 10% butanone for 1h (Fig. 7a). We performed chemotaxis assays by evaluating attraction toward butanone (normally a weak attractant) vs. isoamyl alcohol (a natural strong attractant) for naïve and conditioned animals. Naïve animals (i.e. non-conditioned) are attracted almost exclusively to isoamyl alcohol whereas a significant proportion of food-butanone conditioned animals choose butanone over isoamyl alcohol (Fig. 7a). This increase in chemotaxis toward butanone of conditioned animals relative to naïve animals corresponds to the learning index.

**Figure 7.**
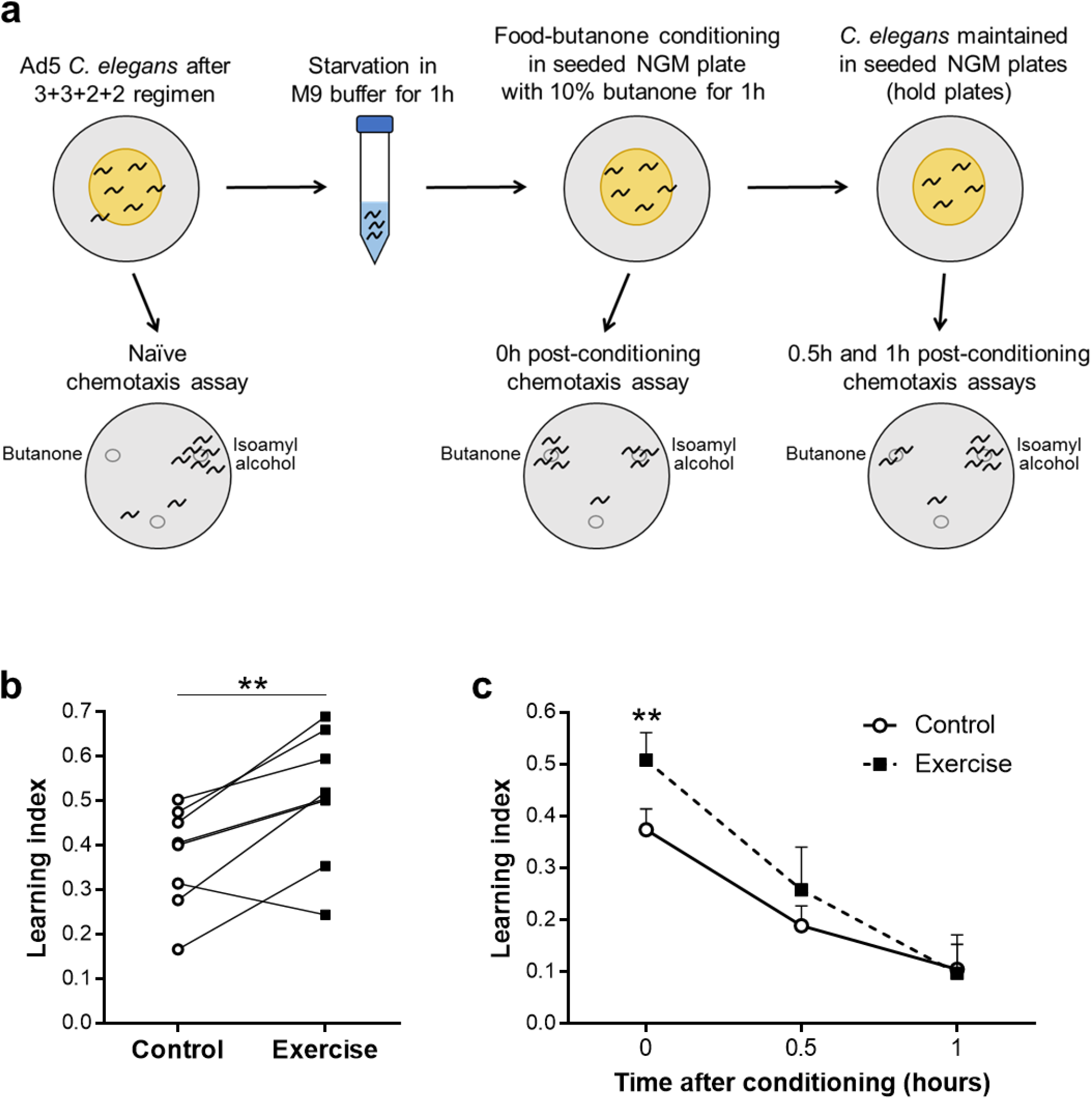
Learning ability is enhanced in exercised *C. elegans*. **a**, Diagram of the associative learning assay. After the 3+3+2+2 regimen, we starved Ad5 animals in M9 buffer for 1h followed by food-butanone conditioning in a seeded NGM plate with 10% butanone solution on the inside of the lid. After conditioning, we maintained animals in seeded NGM plates (hold plates). At different points of the protocol, we used part of the nematode population for naïve and post-conditioning chemotaxis assays. We performed chemotaxis assays by testing animal attraction to 10% butanone vs. 1:1000 isoamyl alcohol. Naïve animals are attracted almost exclusively to isoamyl alcohol whereas a significant proportion of food-butanone conditioned animals choose butanone over isoamyl alcohol. **b**, Learning index of Ad5 WT animals 0h post-conditioning after exposure to the 3+3+2+2 regimen. Chemotaxis index (CI) = (animal number at butanone – animal number at isoamyl alcohol)/(total animal number – immobile animal number at origin). We calculated learning index by subtraction of naïve CI from post-conditioning CI. *n* = 8 independent trials. We used 200-400 animals per sample in each trial. Statistical significance determined by paired two-tailed Student’s *t* test. **c**, Learning index of Ad5 WT animals for up to 1h post-conditioning after exposure to the 3+3+2+2 regimen. Number of independent trials performed: *n*_0h_ = 8, *n*_0.5h_ = 4, *n*_1h_ = 4. We used 200-400 animals per sample in each trial. Statistical significance determined by paired two-tailed Student’s *t* test. Data are mean ± s.e.m. \*\**P*<0.01.

Remarkably, we find that the 3+3+2+2 regimen increased the learning index of Ad5 exercised animals by an average of 35% as compared to non-exercised control counterparts (Fig. 7b). Despite the quantifiable increase in learning ability, short-term memory of exercised animals did not significantly improve – both control and exercised animals showed a 50% reduction in learning index 0.5h post-conditioning and just a residual learning index 1h post-conditioning (Fig. 7c). These results reveal that long-term swim exercise enhances neuronal learning ability in adult *C. elegans* and raise the question as to whether specific neuronal circuits may be aided by particular exercise experiences.

### Long-term swim exercise improves neuronal health in multiple *C. elegans* neurodegeneration models

An additional measure of neuronal health is the capacity to respond to stress conditions induced by neurotoxic proteins. We therefore sought to determine whether exercise could counter pathological conditions in *C. elegans* neurodegenerative disease models. We started by analyzing a *C. elegans* strain that expresses the human aggregating F3ΔK280 Tau fragment together with the full-length (FL) mutant Tau V337M in all neurons (*P_rab-3_F3ΔK280; P_aex-3_h4R1NTauV337M*) [50]. This tauopathy model is based on mutations commonly identified in patients with frontotemporal dementia with Parkinsonism linked to chromosome 17 (FTDP-17) [51]. Transgenic strains exhibit robust toxicity phenotypes, including strongly impaired motility, reflected in a reduced mean velocity compared to anti-aggregating strains [50]. As shown in Fig. 8a, the 3+3+2+2 swim regimen led to a clear increase in mean velocity in exercised aggregating Tau animals, at Ad5, Ad8, and Ad11, relative to non-exercised control counterparts. We conclude that swim exercise can protect against the deleterious consequences of aggregating Tau.

**Figure 8.**
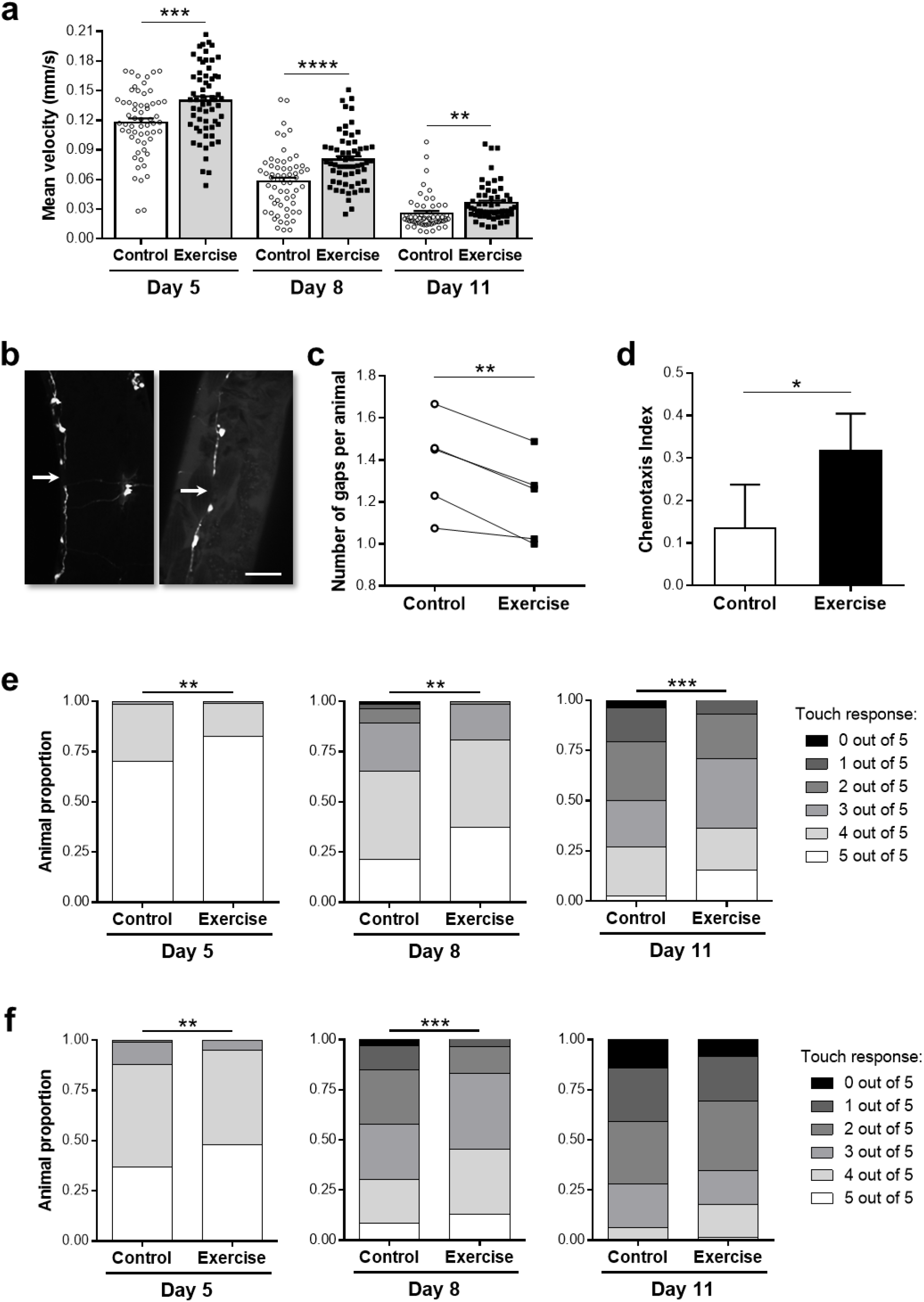
Long-term swim exercise improves neuronal health in multiple *C. elegans* neurodegeneration models. **a**, Mean velocity of Ad5, Ad8, and Ad11 animals expressing aggregating Tau in all neurons (*P_rab-3_F3ΔK280; P_aex-3_h4R1NTauV337M*) exposed to the 3+3+2+2 regimen. We measured mean velocity of single animals placed on an unseeded NGM plate over a 30-second time period. Note that Tau-expressing animals did not exhibit a standard swimming motion due to impaired motility but were more active in liquid than on agar so enhanced activity is confirmed. Each point represents a single animal. Number of animals used for analysis: *n*_Ad5 Control_ = 59, *n*_Ad5 Exercise_ = 60, *n*_Ad8 Control_ = 60, *n*_Ad8 Exercise_ = 60, *n*_Ad11 Control_ = 54, *n*_Ad11 Exercise_ = 57. Data compiled from 3 independent trials. Statistical significance determined by unpaired two-tailed Student’s *t* test. **b**, Representative confocal images of Ad3 animals expressing aggregating Tau in all neurons and GFP in GABAergic motor neurons (*P_rab-3_F3ΔK280; P_aex-3_h4R1NTauV337M; P_unc-25_GFP*). Arrows indicate gaps in the ventral cords. Scale bar: 30 µm. **c**, Average number of gaps detectable in ventral and dorsal cords of Ad3 animals expressing aggregating Tau in all neurons and GFP in GABAergic motor neurons exposed to the 3+2 regimen. Note that this strain did not exhibit a standard swimming motion due to the severe uncoordinated phenotype, but animals were still more active in liquid than on agar so enhanced activity is confirmed. Quantification of gaps in nerve cords was performed blind to exercise condition. *n* = 5 independent trials. We used 30-40 animals per sample in each trial. Statistical significance determined by paired two-tailed Student’s *t* test. **d**, Chemotaxis index toward benzaldehyde of Ad3 animals expressing neuronal Aβ_1-42_ (*smg-1(cc546ts) P_snb-1_Aβ_1-42_::long 3’-UTR*) exposed to the 3+3 regimen. We raised animals at 23°C from the egg stage onward. For the chemotaxis assay, we divided 60 mm unseeded NGM plates in two halves and spotted 0.1% benzaldehyde at the edge of one half and absolute ethanol at the edge of the other half. The chemotaxis index (CI) was calculated as follows: CI = (animal number at benzaldehyde half – animal number at ethanol half)/(total animal number – immobile animal number at origin). *n* = 5 independent trials. We used 40-50 animals per sample in each trial. Statistical significance determined by paired two-tailed Student’s *t* test. **e, f**, Anterior (**e**) and posterior (**f**) touch sensitivity of Ad5, Ad8, and Ad11 animals expressing polyQ128 in the touch receptor neurons (*P_mec-3_htt57Q128*) exposed to the 3+3+2+2 regimen. Animals were scored blind to exercise condition. Number of animals used for analysis in **e** and **f**: *n*_Ad5 Control_ = 248, *n*_Ad5 Exercise_ = 233, *n*_Ad8 Control_ = 155, *n*_Ad8 Exercise_ = 147, *n*_Ad11 Control_ = 122, *n*_Ad11 Exercise_= 119. Data compiled from 5 independent trials. Statistical significance determined by chi-square test. Data are mean ± s.e.m. \**P*<0.05, \*\**P*<0.01, \*\*\**P*<0.001, \*\*\*\**P*<0.0001.

A second consequence of aggregating tau expression in the *C. elegans* nervous system are severe morphological abnormalities in nerve cords, such that gaps in the ventral and dorsal cords can be observed during developmental and adult stages when GFP reporters are used for their visualization [50]. We used a *C. elegans* strain in which the GABAergic motor neurons are GFP-labeled in the aggregating Tau background (*P_rab-3_F3ΔK280; P_aex-3_h4R1NTauV337M; P_unc-25_GFP*) [50] to quantify the number of gaps in GABAergic motor neurons in exercised vs. non-exercised control animals (Fig. 8b). We found that this strain exhibited severely uncoordinated locomotion; therefore, we elected to reduce the swim regimen to the first two days of adulthood (3+2 regimen), with analysis of cord morphology one day later. We find that swim-exercised Ad3 mutants exhibited a subtle but consistent decrease in the average number of gaps in GABAergic motor neurons relative to non-exercised controls (Fig. 8c). Taken together, our analysis of a tauopathy model demonstrates that swim exercise is able to improve pathological effects of neuronal aggregating Tau, both at the functional and morphological levels.

We also addressed impact of exercise on a *C. elegans* AD model in which human amyloid-β (Aβ) peptide (1-42) is pan-neuronally expressed in a temperature-sensitive manner (*smg-1(cc546ts) P_snb-1_Aβ_1-42_::long 3’-UTR*) [52]. One of the phenotypes previously described in this strain is a reduced chemotactic ability toward benzaldehyde [52]. When raised at 23°C (a temperature that induces expression of Aβ_1-42_), chemotaxis performance of transgenic Aβ animals decreases rapidly during the first days of adulthood. We decided to limit the swim regimen for this strain to the first two days of adulthood (3+3 regimen) with assay at Ad3 because of the severity of chemotactic deficits exhibited by non-exercised animals. We find that exercised Ad3 Aβ_1-42_-expressing animals exhibited a significantly higher chemotaxis index toward benzaldehyde than non-exercised control counterparts (Fig. 8d). Thus, swim exercise improves a neuronally controlled behavior in a *C. elegans* model of AD-associated toxicity.

Finally, we tested exercise effects in a *C. elegans* HD model in which the first 57 amino acids of human Huntingtin protein (HTT) are fused to an expanded polyglutamine (polyQ128), and this transgene is expressed in ten neurons including the six touch receptor neurons (*P_mec-3_htt57Q128*) [53]. The *C. elegans* touch receptor neurons mediate the response to gentle touch [54] and polyQ expression in these specific neurons greatly accelerates touch deficits during adult life [53, 55]. After a 3+3+2+2 swim regimen, we performed touch sensitivity assays by quantifying the animal responsiveness to five gentle touches on the head or the tail at different days of adulthood. In studies in which investigators were blinded to previous exercise experience of the animals, we find that exercised polyQ animals exhibited a significant improvement in touch sensitivity at Ad5, Ad8, and Ad11 for the anterior region (Fig. 8e) and at Ad5 and Ad8 for the posterior region (Fig. 8f). Taking advantage of touch sensitivity as a direct readout of touch receptor neuron function, we show that long-term swim exercise in *C. elegans* improves neuronal healthspan in a sensitized polyQ-mediated toxicity model.

Overall, our testing of exercise impact on models in which human disease proteins impair *C. elegans* adult neuron functions demonstrates that long-term swim exercise improves neuronal healthspan at both morphological and functional levels, for different neuronal cell types in multiple neurodegeneration models. A particularly striking corollary is that impact of sustained early adult swim training can confer a demonstrable health benefit even long after training has ended.

## Discussion

The development of short-lived genetic models in which fundamental questions of exercise biology can be evaluated during the entire aging process holds great potential for advancing our knowledge on trans-tissue exercise signaling that improves old-age health. Here, we define long-term adult exercise regimens that induce robust exercise adaptations in multiple *C. elegans* tissues. We show that repeated daily swim sessions are essential for molecular, morphological, and functional adaptations at the body wall muscle level. Moreover, whole-animal health parameters, including mitochondrial respiration and mid-life survival, are improved by swim exercise. An important point is that we find that long-term exercise extends functionality of tissues not directly involved in physical activity, such as pharynx and intestine. Furthermore, we show that swim exercise improves learning ability of WT animals and is able to mitigate the consequences of neuronal pathologies in tauopathy, AD, and HD *C. elegans* models. Particularly noteworthy is that benefits of exercise regimens in young adult life can be documented to extend late into life, which suggests a potential switch mechanism for healthy aging might be triggered by early enhanced activity.

### How *C. elegans* trains for sustained adult health

In search of the ideal exercise regimen, we found that multiple swim sessions per day, over several days, are essential for exercise-dependent molecular and functional adaptations in *C. elegans*. This finding reveals that highly regular exercise bouts are key for promoting the cumulative effects of single exercise sessions in *C. elegans*. Using muscle gene expression and locomotion performance as readouts, we identified the 3+3+2+2 regimen as the most consistent protocol to induce long-term exercise adaptations. Other exercise regimens, however, also promoted relevant changes; for example, the 4+4+4+4 regimen induced the best locomotion performance at Ad5 and clear mitochondrial adaptations, together with a mixed muscle transcriptional response. Thus, it is possible that particular health and fitness readouts might require a slightly adjusted multiple daily swim regimen to induce optimal exercise adaptation for a given parameter. Still, that the 3+3+2+2 protocol confers a broad swath of health improvements sets the stage for detailed mechanistic dissection of the molecular circuits that maintain animal vigor as a consequence of exercise.

Our data establishing the need for repeated exercise as required for health adaptations may partially explain the lack of mitochondrial adaptations and survival improvement previously reported for a vibration *C. elegans* exercise model [19] and the lack of locomotion improvement in a Duchenne muscular dystrophy (DMD) *C. elegans* model swim-exercised 90 min per day [20], as these studies did not include exercise training in consecutive days or multiple times per day.

In addition, when working with mutant or transgenic strains with severe health deficits or aging animals, our data indicate that the exercise regimen should be reduced to prevent overtraining and consequent deleterious effects. Similar adaptations are routinely made in human physical therapy regimens. The transient increase in physical activity during an exercise bout and the following post-exercise recovery period are crucial for long-lasting exercise adaptation in mammals [29, 56, 57]. We introduced a 4.5h minimum interval between swim sessions in our exercise regimens based on our previous documentation that most acute exercise changes return to baseline levels by 4h post-exercise [15]. As rest and recovery appears to be important to instituting positive outcomes of exercise, we suggest that recently described regimens in *C. elegans* based on continuous burrowing or swimming over several days [20] may need to incorporate activity breaks in order to accomplish long-term benefits.

### Strong muscle and healthy mitochondria into old age

Body wall muscle aging in *C. elegans* is characterized by a marked downregulation of muscle structural genes followed by a decline in mitochondrial network organization, which strongly correlates with loss of movement [26, 31]. Our swim exercise regimens are able to delay sequential muscle aging processes, resulting in increased locomotor performances in 2D (crawling) and 3D (burrowing) environments. Mitochondrial improvements likely extend beyond body wall muscle since we found that whole-animal mitochondrial respiration parameters were significantly changed after long-term swim exercise. The improvement in mitochondrial efficiency and health was apparent in exercised animals from the increase in basal respiration, maximal respiration, spare capacity, and ATP-linked respiration, and from the reduction in proton leak. The exercise-dependent improvement in multiple respiration parameters was more pronounced than previously reported by Hartman et al. [16], which might be explained by the different swim exercise regimens used in that study and ours.

### Early life exercise can extend median lifespan of an active population

A majority of human exercise studies support that exercise can reduce mortality risk and increase longevity of healthy subjects, patients, and elite athletes, although some studies suggest a negative or neutral association between exercise and lifespan in humans [35–38]. Thus, a fair generalization may be that the extent of the exercise benefits is variable and dependent in part on the type of exercise, the population studied, and the particular outcomes recorded. Similar to observations in humans, the effect of exercise on *C. elegans* longevity seems to be highly dependent on the adopted exercise regimen with reported outcomes in previous studies varying from detrimental to beneficial [16–20]. Our swim exercise regimen promoted a consistent increase in mid-life population survival rather than an increase in maximum lifespan. Thus, exercised populations are able to survive extra days during the time-window when the mortality rate is at its peak. It is important to note that in our studies, this time-window occurs 8 to 16 days after the last exercise session, raising the possibility that extension of the swim regimen beyond the first four days of adulthood may further potentiate survival differences and underscoring that early adult activity can have long lasting impact on healthy aging (see further discussion below).

### Systemic benefits of exercise

One of the pressing questions in exercise biology is how physical activity can promote health benefits in multiple tissues of the body, especially in those tissues not directly engaged by neuromuscular activity. With this issue in mind, we explored whether *C. elegans* exercise promotes health benefits in tissues other than body wall muscle. We found that long-term swim exercise in *C. elegans* increases pharyngeal pumping rate during mid-life, reflecting a delay in the age-associated decline of cardiac-like muscle function [39]. In the case of *C. elegans*, the pumping mechanism is inhibited rather than accelerated during swim training [40], so pharyngeal muscle itself does not “train” – yet pharyngeal functional health nonetheless improves with exercise. Our observations raise the possibility that some exercise benefits to human heart might be conferred via trans-tissue signals rather than exercise-enhanced heart contraction.

We also observed a significant improvement of the older age intestinal barrier function in exercised *C. elegans*, establishing that enhanced gut health is an “off-target” effect of exercise experience. In humans, exercise has been shown to reduce the risk of gastrointestinal cancer, diverticulitis, and inflammatory bowel disease [58] (although high-intensity, prolonged exercise sessions will cause an acute increase in intestinal permeability (or ‘leaky gut’) [59], which has been associated with infectious diseases, obesity, and autoimmune diseases [60]). We suggest that an exercise-dependent hormetic effect on intestinal function can reconcile these apparently contradictory findings – acute exercise bouts may enhance intestinal permeability, but regular exercise training will promote intestinal adaptation over time that might enhance intestinal barrier function, similar to what we detected in *C. elegans*.

Our pharyngeal- and intestinal-related results also highlight the importance of selecting the right time points for the assay of long-term exercise outcomes, since exercise-dependent benefits become apparent at different stages of *C. elegans* adult life depending on the tested tissue. Differences are possibly best assayed soon after the point of known age-associated decline, where the window of health will initially extend. Regardless, the documentation of trans-tissue impact of exercise makes it clear that *C. elegans* can be used as a model for the future molecular dissection of the genetic factors that translate exercise into whole-animal benefit.

### Physical activity is markedly protective for age-associated neuronal decline and neurodegeneration stresses

Our study presents the first clear evidence that exercise in *C. elegans* promotes neuronal health at multiple levels. Kauffman et al. have shown that the learning process is molecularly distinct from memory formation; and that capacity for learning declines with age [13]. We show that learning ability is markedly increased in swim-exercised WT animals, whereas short-term memory retention is not significantly improved. Interestingly, other longevity paradigms have been reported to have focused impact on particular components of the learning/memory process [13] – dietary restricted *eat-2* mutants retain the ability to learn for longer and maintain short- and long-term memory with age, but *daf-2* reduced insulin signaling mutants retain the ability to learn for longer but lose long-term memory at the same rate as WT [13]. Overall, the fact that swim exercise enhances adult learning capacity reveals that particular neuronal capabilities can be improved as a result of animal activity and adds maintained neuronal function to the list of healthspan benefits of exercise in *C. elegans*.

Numerous studies in animal models and humans show that exercise can improve cognitive function and reduce the risk of developing dementia later in life [45–48]. However, the effect of exercise on dementia patients is not clear, with outcomes of published studies ranging from detrimental to beneficial effects of exercise [61–65]. Taking advantage of neurodegeneration models of human disease in *C. elegans*, we show that swim exercise improves neuronal healthspan in disease protein-stressed neurons, at both morphological and functional levels, for different neuronal cell types. We detected exercise-dependent health benefits for motor neurons, chemosensory neurons, and touch receptor neurons in tauopathy, AD, and HD conditions, respectively, revealing a broad impact of exercise in maintaining health in the *C. elegans* nervous system confronted with disease stresses. Exercise prescription by primary care physicians is currently recommended as a way to improve patients’ general health and reduce all-cause mortality [66, 67]. Our data strengthens the hypothesis that exercise can have powerful beneficial effects in neurodegenerative disease patients.

### Exercise during early adult life can have clear long-term impact on later adult health

We show in this study that long-term swim exercise in *C. elegans* induces health benefits at the body wall muscle, pharyngeal, intestinal, and neuronal levels. Remarkably, most of the exercise-dependent benefits are long lasting, with effects still detectable up to 7-16 days after the last exercise session for specific assays. Interestingly, a recent study shows that human volunteers who enrolled in an 8-month exercise regimen still exhibit improvements in aerobic capacity and metabolic health 10 years after completion of exercise training [68]. Thus, persistent health benefits of previous exercise training may be conserved from nematodes to humans. If so, the establishment of *C. elegans* as an exercise model opens the exciting possibility of uncovering the molecular signals responsible for life-long, exercise-dependent systemic health benefits.

## Materials and Methods

### *C. elegans* strains and maintenance

The *C. elegans* strains used in this study were: N2 (WT strain), SJ4103 *zcIs14[P_myo-3_mitoGFP]* [69], BR6563 *byIs161[P_rab-3_F3ΔK280; P_myo-2_mCherry]; bkIs10[P_aex-3_h4R1NTauV337M; P_myo-2_GFP]* [50], BR5707 *byIs161[P_rab-3_F3ΔK280; P_myo-2_mCherry]; bkIs10[P_aex-3_h4R1NTauV337M; P_myo-2_GFP]; juIs73[P_unc-25_GFP] III* [50], CL2355 *smg-1(cc546ts) dvIs50[P_snb-1_Aβ_1-42_::long 3’-UTR P_mtl-2_GFP] I* [52], and ID1 *igIs1[P_mec-3_htt57Q128::CFP, P_mec-7_YFP, lin-15(+)]* [53]. We maintained nematodes at 20°C on NGM plates seeded with live *Escherichia coli* OP50-1 (streptomycin-resistant strain) as previously described [70], unless otherwise stated.

### Long-term swim exercise protocols

We carefully maintained *C. elegans* at 20°C for several generations without being starved prior to any exercise experiment. We used two different methods for our long-term swim exercise experiments: the picking method and the washing method. We performed the picking method for all the experiments presented in this study, except for the oxygen consumption rates and associative learning assays in which the washing method was used due to the high number of animals required for each assay.

For the picking method, we synchronized the population by allowing 30-40 gravid hermaphrodites to lay eggs in a 60 mm seeded NGM plate for 3h before removing the parents. After 64-68h, we began the long-term exercise regimens by picking approximately 60 Ad1 animals to a 60 mm unseeded NGM plate (non-exercise control) or to a 35 mm unseeded NGM plate flooded with 1.5 mL of M9 buffer (exercise). After 90 min, animals from both conditions were picked to 60 mm seeded NGM plates. We repeated this procedure multiple times over the first four days of adulthood depending on the exercise regimen.

For the washing method, we synchronized the population by allowing 300-400 gravid hermaphrodites to lay eggs in a 100 mm seeded NGM plate for 3h before removing the parents. After 64-68h, we began the long-term exercise regimens by washing the Ad1 animals off the plate with M9 buffer into a 15 mL conical tube and allowing them to gravity settle (washing step performed by a gentle swirl of the plate to minimize bacterial transfer). Then, using a glass Pasteur pipette to minimize nematode loss, we split the pellet between a 100 mm unseeded NGM plate (non-exercise control) and a 100 mm unseeded NGM plate flooded with 8-10 mL of M9 buffer (exercise). After 90 min, animals from both conditions were washed off with M9 buffer into 15 mL conical tubes and allowed to settle under gravity. We transferred animals with a glass Pasteur pipette to 100 mm seeded NGM plates. We repeated this washing method multiple times over the first four days of adulthood depending on the exercise regimen.

We performed swim exercise sessions at the following timings: 1+1+1+1 regimen, 1:00 PM each day; 2+2+2+2 regimen, 7:00 AM and 1:00 PM each day; 3+3+2+2 regimen, 7:00 AM, 1:00 PM, and 7:00 PM on the first two days, 7:00 AM and 7:00 PM on the last two days; 3+3+3+3 regimen, 7:00 AM, 1:00 PM, and 7:00 PM each day; 4+4+4+4 regimen, 7:00 AM, 1:00 PM, 7:00 PM, and 1:00 AM each day.

### RNA extraction and quantitative PCR

We performed RNA extraction and qPCR as previously described [15]. Briefly, we collected WT *C. elegans* (20-40 animals per sample) into TRIzol Reagent (Thermo Fisher Scientific) and immediately froze animals in liquid nitrogen. After three freeze-thaw cycles, we extracted total RNA following the manufacturer’s instructions (Thermo Fisher Scientific) and synthesized cDNA from 400 ng of total RNA using the SuperScript III First-Strand Synthesis System (Thermo Fisher Scientific). We carried out qPCR using diluted cDNA, PerfeCTa SYBR Green FastMix (Quantabio), and 0.5 μM of gene-specific primers (Table S2) in a 7500 Fast Real-Time PCR System (Thermo Fisher Scientific), using a two-step cycling protocol (3 seconds at 95°C, 30 seconds at 60°C) for 40 cycles. We calculated relative expression using the ΔΔCt method with *cdc-42* and Y45F10D.4 as reference genes.

### Maximum and mean velocity

We measured maximum and mean velocity as previously described [10], with minor modifications. Briefly, we transferred single animals to a 60 mm unseeded NGM plate and immediately recorded a 30-second video at a rate of frames per second with a Prosilica GC 1380 camera (Allied Vision) connected to a Macro Zoom 7000 lens (Navitar) using StreamPix 5 software (NorPix). We used approximately 20 animals per trial. After recording, we returned animals to a 60 mm seeded NGM plate for re-use in later time points. Maximum and mean velocity were calculated with the ImageJ plugin wrMTrck [71].

### Microfluidic assays

We synchronized the population by allowing 30-40 gravid hermaphrodites to lay eggs in a 60 mm seeded NGM plate for 3h before removing the parents. After 64-68h, we began the 3+3+2+2 regimen by inserting approximately 80 Ad1 animals (5-15 animals per chamber) in small chamber height microfluidics (non-exercise control) and in large chamber height microfluidics (exercise), both filled with M9 buffer. After 90 min, animals from both conditions were washed off the microfluidic chambers with M9 buffer and returned to 60 mm seeded NGM plates. We repeated this procedure for a total of 10 times over the first four days of adulthood at the standard timings for the 3+3+2+2 regimen. We used microfluidic chambers with the following heights: 20 µm for Ad1 control animals, 30 µm for Ad2-4 control animals, and ~300 µm for all exercise animals. Representative videos were recorded at a rate of 20 frames per second with a Prosilica GC 1380 camera (Allied Vision) connected to a Macro Zoom 7000 lens (Navitar) using StreamPix 5 software (NorPix).

### Burrowing Assay

We prepared a 26% (w/v) Pluronic F-127 (Sigma-Aldrich) solution in water several days before the assay and stored it at 4°C to facilitate complete dissolving. The Pluronic F-127 solution remains liquid at cooler temperatures, but fully gels to a solid at room temperature. At least one hour before the burrowing assay, we placed the Pluronic F-127 solution at 14°C and allowed it to equilibrate before use (at this temperature it remains fully liquid). We carried out the burrowing assay at room temperature (20°C) by adding 30 µL of Pluronic F-127 solution to the center of a well in a 12-well plate. We immediately transferred approximately 40-45 animals to the 30 µL droplet using a nematode pick. Once the droplet was fully gelled, we added ~4 mL of additional Pluronic F-127 solution to form a final gel height of ~0.75 cm. After the top layer of Pluronic F-127 had fully gelled, we added 20 µL of *E. coli* OP50-1 chemoattractant solution (100 mg of *E. coli* OP50-1 per mL of liquid NGM) to the center of the gel surface (this was considered t = 0 min). The number of animals that reached the gel surface was scored at regular intervals up to 3h. For each trial, we performed the burrowing assay in triplicate.

### Confocal microscopy

We fixed *C. elegans* in 2% paraformaldehyde/phosphate buffered saline for 30 min at room temperature and then stored them at 4°C in M9 buffer until imaging. We performed confocal imaging with a CSU-X1 spinning disk unit (Yokogawa) mounted to an Axio Imager Z1 microscope (ZEISS) using MetaMorph Premier software (Molecular Devices). For *P_myo-3_mitoGFP* animals (SJ4103 strain), we imaged two different regions of the anterior body wall muscle per animal, and scored images blind to exercise condition using Blinder software [33]. For the animals with GFP-labeled GABAergic motor neurons in the pro-aggregant Tau background (BR5707 strain), we used 30-40 animals per trial and quantified the total number of gaps in the ventral and dorsal cords per animal. Quantification was performed blind to exercise condition.

### Oxygen consumption rates

We exercised animals with the washing method using the 4+4+4+4 regimen because at the time of these experiments we had gathered preliminary data that suggested the 4+4+4+4 regimen as one of the best to promote long-term exercise adaptation (later optimization revealed the 3+3+2+2 regimen as the most consistent exercise protocol).

We measured oxygen consumption rates as previously described [16], with some modifications. We washed animals off the plate with M9 buffer into a 15 mL conical tube. After settling by gravity, animals were transferred to a 100 mm unseeded NGM plate for 20 min to get rid of bacteria. We picked 20-25 animals to each well of a Seahorse XF24 Cell Culture Microplate (Agilent) with 150 µL of M9 buffer, leaving 2-4 wells for blanks. We added 375 µL of EPA water (a lower-saline solution to ensure solubility of drugs [34]) to each well. All oxygen consumption rate measurements were performed in a Seahorse XFe24 Analyzer (Agilent). Basal respiration was measured first in every well followed by injection of Carbonyl cyanide-p-trifluoromethoxyphenylhydrazone (FCCP) in half of the wells to uncouple mitochondria or dicyclohexylcarbodiimide (DCCD) in the other half of the wells to inhibit ATP synthesis. Sodium azide was injected at the end of the assay in every well to completely block mitochondrial respiration. All respiration parameters were normalized to the number of animals per well.

### Lifespan assay

After the 3+3+2+2 regimen, we started a lifespan assay with approximately 40-45 animals per 60 mm NGM plate. We scored and moved animals to fresh plates daily until Ad8 and every two days after that. Animals that bagged or crawled off the plates were censored. Animals were considered dead when they mounted no response to vigorous prodding.

Exercised animals lived longer than non-exercised controls between Ad12 and Ad20. During this period, the percentage survival increase for the exercise population ranged from ~4% on Ad12 to ~20% on Ad16. We calculated the average survival increase for exercised animals using Ad12, Ad14, Ad16, Ad18, and Ad20 values.

### Pharyngeal pumping rate

We recorded 30-second videos at a rate of 30 frames per second of *C. elegans* on seeded NGM with a Lt225 camera (Lumenera) connected to an Axiovert 35M microscope (ZEISS) using StreamPix 6 software (NorPix). We used approximately 25 animals per trial. We counted the number of pharyngeal pumps during the 30 seconds manually in ImageJ.

### Smurf assay

We performed Smurf assays as previously described [44], with minor modifications. Briefly, 25-40 animals were incubated for 3h at 20°C in 5% (w/v) of blue food dye (Erioglaucine disodium salt, Sigma-Aldrich) in *E. coli* OP50-1 liquid culture grown overnight in lysogeny broth (LB). We then transferred the animals to an unseeded NGM plate and immobilized them with 1M sodium azide before scoring for intestinal leakage in a SZX2 microscope (Olympus). Representative images were taken with a 14MP MU1403 microscope digital camera (AmScope) using AmScope 3.7 software (AmScope). Animals with blue food dye in the germline were not considered Smurf animals unless the dye was also observed in the body cavity.

### Associative learning assay

We performed associative learning assays as previously described [13, 72], with some modifications. We exercised animals with the washing method and, on the day of the assay, we washed animals off the plate with M9 buffer into a 15 mL conical tube. After three washes with M9 buffer, we used some animals for naïve chemotaxis assay and starved the rest in 4 mL of M9 buffer for 1h. After starvation, animals were conditioned in a 100 mm seeded NGM plate with three 2 µL streaks of 10% butanone (Sigma-Aldrich) in absolute ethanol placed on the inside of the lid. We parafilmed the plate and incubated it for 1h at room temperature. We washed animals off the plate with M9 buffer into a 15 mL conical tube. After two washes with M9 buffer, we used some animals for the 0h post-conditioning chemotaxis assay and split the rest between two 100 mm seeded NGM plates (hold plates). After 0.5h or 1h at room temperature, we washed animals off the hold plates with M9 buffer into a 15 mL conical tube, washed them two additional times with M9 buffer, and used them for the 0.5h and 1h post-conditioning chemotaxis assays.

For the chemotaxis assays, we marked 100 mm unseeded NGM plates with spots on the bottom (origin) and each side of the plate [72]. We spotted 1 µL of 10% butanone in absolute ethanol at one side of the plate and 1 µL of 1:1000 isoamyl alcohol (Sigma-Aldrich) in absolute ethanol at the other side of the plate. We also spotted 1 µL of 1M sodium azide at each side of the plate to paralyze animals that reached the odorant spots. Approximately 100-200 animals were transferred with a glass Pasteur pipette to the origin spot on each plate, and incubated for 1h at room temperature. We performed two technical replicates for each chemotaxis assay. Images were taken with a Prosilica GC 1380 camera (Allied Vision) connected to a Macro Zoom 7000 lens (Navitar) using StreamPix 5 software (NorPix) at the start of the assay to count the total number of animals and at the end of the assay to count the number of animals at each spot. Chemotaxis index (CI) was calculated as follows: CI = (animal number at butanone – animal number at isoamyl alcohol)/(total animal number – immobile animal number at origin). Learning index (LI) was calculated as follows: LI = CI_Post-conditioning_ – CI_Naïve_.

We found it essential to use isoamyl alcohol for the chemotaxis assays because Ad5 naïve animals were strongly attracted to butanone when absolute ethanol was used as the control odorant. Conversely, Ad5 naïve animals were attracted almost exclusively to isoamyl alcohol rather than butanone, which vastly increased the dynamic range over which to test how exercise affected the food-butanone association.

### Chemotaxis assay toward benzaldehyde

We maintained the CL2355 strain, which enables temperature-sensitive expression of neuronal Aβ_1-42_, at 16°C. We raised animals at 23°C (i.e. expression of Aβ_1-42_ induced) from the egg stage onward for the exercise experiments. For the chemotaxis assay, we divided 60 mm unseeded NGM plates in two halves and spotted 1 µL of 0.1% benzaldehyde (Sigma-Aldrich) in absolute ethanol at the edge of one half and 1 µL of absolute ethanol at the edge of the other half. After moving animals to an unseeded NGM plate to get rid of bacteria, we transferred 20-25 animals to the center of the chemotaxis plate and incubated for 2h at 23°C. We performed two technical replicates for each chemotaxis assay. Chemotaxis index (CI) was calculated as follows: CI = (animal number at benzaldehyde half – animal number at ethanol half)/(total animal number – immobile animal number at origin).

### Touch sensitivity assay

We performed touch sensitivity assays as previously described [73], with minor modifications. Briefly, animals were scored blind to exercise condition by a gentle touch with a thin platinum wire (0.025 mm of diameter, Alfa Aesar) in the anterior region followed by a gentle touch in the posterior region a few seconds later. We repeated this procedure for a total of 5 anterior and 5 posterior touches per animal. Normally, a positive touch response resulted in animals backing away from the touch.

### Statistical analyses

The data in this study are presented as the mean ± standard error of the mean (s.e.m.). We performed a minimum of three independent trials for each experiment. The specific number of data points and the test used for each statistical analysis are presented in the figure legends.

We used log-rank test to compare the proportion over time of control vs. exercise populations that reached a specific phenotype (burrowing and lifespan assays). We used chi-square test to compare the distribution into multiple categories of control vs. exercise populations (muscle mitochondrial morphology and touch sensitivity assays). For all other assays, we used two-tailed Student’s *t* test to determine statistical significance given the comparison of control vs. exercise means for a single time point. Paired Student’s *t* test was used when control and exercise values consisted of matched pairs, whereas unpaired Student’s *t* test was used when control and exercise values were compiled from multiple trials.

## Acknowledgements

We thank the following people for providing *C. elegans* strains: Ralf Baumeister for BR6563 and BR5707, and J. Alex Parker and Christian Néri for ID1. The other strains used in this study were provided by the *Caenorhabditis* Genetics Center (CGC), which is funded by NIH Office of Research Infrastructure Programs (P40 OD010440). We thank Leila Lesanpezeshki for help with production of microfluidic devices and Charline Borghgraef for help with protocol optimization of associative learning assays.

