## Supplementary material for "Swim exercise in *C. elegans* extends neuromuscular and intestinal healthspan, enhances learning ability, and protects against neurodegeneration": Table S1

**Table S1.** Genes selected for qPCR analysis.

| ***C. elegans*** **genes** | **Mammalian homologs** |
|---|---|
| *mup-2* | Troponin T |
| *tnt-2* | Troponin T |
| *unc-27* | Troponin I |
| *lev-11* | Tropomyosin 1 |
| *unc-15* | Paramyosin |
| *mlc-1* | Myosin light chains |
| *unc-54* | Myosin heavy chains |
| *myo-3* | Myosin heavy chains |
| *zig-12* | Titin |
| *unc-87* | Calponin-like |
