## Supplementary material for "Swim exercise in *C. elegans* extends neuromuscular and intestinal healthspan, enhances learning ability, and protects against neurodegeneration": Table S2

**Table S2.** Primers used for qPCR.

| Gene | Forward primer (5'-3') | Reverse primer (5'-3') | Amplification efficiency |
|---|---|---|---|
| ***cdc-42*** | CTGCTGGACAGGAAGATTACG | CTCGGACATTCTCGAATGAAG | 1.03 |
| ***lev-11*** | CCGCTGAAGAGAAAGTCCGT | TCGTCTCCGGTCTGAGTCAT | 0.93 |
| ***mlc-1*** | TGGAGCCTTTGCCATGTTC | CTTGACCTCATCCTCGTCCAAT | 0.99 |
| ***mup-2*** | AACGCAAGGCTAAGGCTGAT | AGCTCCGGCTTCAACTCTTC | 0.92 |
| ***myo-3*** | AGGGAGACTTGAAGGTTGCG | AGCGAGCTTAGCATTGGTGT | 0.93 |
| ***tnt-2*** | ATGGGGACGCAAAGAGAACG | AATTGGGTTGACTGGTGGCT | 1.00 |
| ***unc-15*** | CGAGGAAGCCAATGGACGTA | AAATCAGCTTGAGCGGTGGA | 0.95 |
| ***unc-27*** | CGTGGAAAGTTCGTCAAGCC | TCTTGAGGTTGGCACGGAAG | 0.98 |
| ***unc-54*** | ACTACCAACACGAAGCCGAG | GGCGTTAGCCTTGGAGAGTT | 0.94 |
| ***unc-87*** | ATGACTGGATTCGGACAGCC | AGCTTGAGAAGCAAAACGGT | 0.98 |
| **Y45F10D.4** | GTCGCTTCAAATCAGTTCAGC | GTTCTTGTCAAGTGATCCGACA | 0.94 |
| ***zig-12*** | GATCAGAGAACGGGTCGGTG | CTCCTCAAGCTCGTCTGGTC | 1.02 |
