## Supplementary material for "Swim exercise in *C. elegans* extends neuromuscular and intestinal healthspan, enhances learning ability, and protects against neurodegeneration": Video legends

**Video S1.** Representative videos of Ad2 WT animals in an exercise-permissive microfluidic chamber (large chamber height) (**a**) and in a control microfluidic chamber (small chamber height non-permissive for swimming) (**b**). Animals are in M9 buffer in both conditions but locomotion changes from swimming (**a**) to crawling (**b**) due to physical confinement in the control chamber.
